# Evaluating the accuracy of DNA stable isotope probing

**DOI:** 10.1101/138719

**Authors:** Nicholas D. Youngblut, Daniel H. Buckley

## Abstract

**Originality-Significance Statement:** By combining DNA Stable Isotope Probing (DNA-SIP) with multiplexed high throughput DNA sequencing (HTS-DNA-SIP), it is now possible to identify patterns of isotope incorporation for thousands of microbial taxa. HTS-DNA-SIP has enormous potential to reveal patterns of carbon and nitrogen exchange within microbial food webs. A current limitation is that, due to the expense of these experiments, it has been impossible to evaluate the accuracy of DNA-SIP methods. We have developed a model that simulates DNA-SIP data, and we use the model to systematically evaluate and validate the accuracy of DNA-SIP analyses. This model can determine the analytical accuracy of DNA-SIP experiments in a range of contexts. Furthermore, the ability to predict experimental outcomes, as a function of experimental design and community characteristics, should be of great use in the design and interpretation DNA-SIP experiments.

**Summary:** DNA Stable isotope probing (DNA-SIP) is a powerful method that identifies *in situ* isotope assimilation by microbial taxa. Combining DNA-SIP with multiplexed high throughput DNA sequencing (HTS-DNA-SIP) creates the potential to map *in situ* assimilation dynamics for thousands of microbial taxonomic units. However, the accuracy of methods for analyzing DNA-SIP data has never been evaluated. We have developed a toolset (SIPSim) for simulating HTS-DNA-SIP datasets and evaluating the accuracy of methods for analyzing HTS-DNA-SIP data. We evaluated two different approaches to analyzing HTS-DNA-SIP data: “high resolution stable isotope probing” (HR-SIP) and “quantitative stable isotope probing” (q-SIP). HR-SIP was highly specific and moderately sensitive, with very few false positives but potential for false negatives. In contrast, q-SIP had fewer false negatives but many false positives. We also found HR-SIP more robust than q-SIP with respect to experimental variance. Furthermore, we found that the detection sensitivity of HTS-DNA-SIP can be increased without compromising specificity by evaluating evidence of isotope incorporation over multiple windows of buoyant density (MW-HR-SIP). SIPSim provides a platform for determining the accuracy of HTS-DNA-SIP methods across a range of experimental parameters, which will be useful in the design, analysis, and validation of DNA-SIP experiments.

## Introduction

Stable isotope probing of nucleic acids (DNA-SIP and RNA-SIP) is a powerful culture-independent method for linking microbial metabolic functioning to taxonomic identity (Radajewski et al., 2003). In particular, DNA-SIP has been used extensively to identify microbial assimilation of various ^13^C- and ^15^N-labeled substrates in a multitude of environments (Uhlík et al., 2009). DNA-SIP identifies microbes that assimilate isotope into their DNA (“incorporators”) by exploiting the increased buoyant density (BD) of isotopically labeled (“heavy”) DNA relative to unlabeled (“light”) DNA. For example, fully ^13^C- and ^15^N-labeled DNA will increase in BD by 0.036 and 0.016 g ml^−1^, respectively (Birnie and Rickwood, 1978).

Ideally, isopycnic centrifugation could be used to completely separate labeled and unlabeled DNA fragments based solely on this difference in BD. However, several factors besides BD can impact the position of DNA in isopycnic gradients. For example, G + C content variation across a single genome can produce unlabeled DNA fragments that vary in BD by up to 0.03 g ml^−1^, while G + C content variation between microbial genomes can cause the average BD of unlabeled DNA fragments to vary by up to 0.05 g ml^−1^ (Youngblut and Buckley, 2014). In addition, DNA in SIP experiments will often be partially labeled as a consequence of isotope dilution from unlabeled endogenous substrates. Therefore, it is unlikely that nucleic acid SIP experiments will ever achieve complete separation of labeled and unlabeled DNA.

In the absence of complete separation between labeled and unlabeled DNA, isotope incorporators must be identified using some statistical procedure suitable for comparing the BD distributions of DNA fragments from labeled and unlabeled samples (Pepe-Ranney et al., 2016a). The use of multiplexed high throughput sequencing with DNA-SIP (“HTS-DNA-SIP”) makes it possible to sequence SSU rRNA amplicons across many density gradient fractions and simultaneously determine the BD distributions for thousands of taxa. The problem then becomes one of identifying those taxa that have increased in BD in the isotopically labeled samples relative to the corresponding unlabeled controls.

Different analytical approaches have been applied to HTS-DNA-SIP datasets to identify changes in DNA BD in response to isotopic labeling. These include “high resolution stable isotope probing” (HR-SIP) and “quantitative stable isotope probing” (q-SIP), which both analyze SSU rRNA amplicons across numerous gradient fractions (Hungate et al., 2015; Pepe-Ranney et al., 2016a; Pepe-Ranney et al., 2016b). However, these methods differ in the statistical procedures used to detect taxa that incorporate isotopic label. HR-SIP identifies isotopically labeled taxa by evaluating the sequence composition of high density “heavy” fractions using a differential abundance quantification framework that evaluates sequence count data in isotopically labeled samples relative to their corresponding unlabeled controls. Differential abundance between the “heavy” fractions of labeled and control gradients is measured with DESeq2 (Love et al., 2014), which uses sophisticated statistical methods to reduce technical error and increase analytical power for analysis of microbiome data (McMurdie and Holmes, 2014). In a very different approach, q-SIP transforms SSU rRNA relative abundance values by using qPCR estimates of total SSU rRNA gene copies present within gradient fractions. These normalized data are used to estimate average BD for each taxon across density gradients for both isotopically labeled samples and corresponding unlabeled controls (Hungate et al., 2015). Incorporators are then determined by using a permutation procedure to identify those taxa whose BD shifts are unlikely to occur as a result of chance.

While DNA-SIP is a powerful method for the discovery and characterization of microorganisms *in situ*, systematic assessment of the specificity or sensitivity of this method has not been performed. Empirical validations of DNA-SIP methods typically include only one or a few organisms (Lueders et al., 2004; Buckley et al., 2007; Cupples et al., 2007; Wawrik et al., 2009; Andeer et al., 2012), and such approaches do not adequately replicate the complexity of the DNA fragment BD distributions expected in a typical DNA-SIP experiment (Youngblut and Buckley, 2014). DNA-SIP experiments vary in the diversity of the target community, DNA G + C content distribution, the number of incorporators, incorporator relative abundance, and the atom % excess of labeled DNA. Systematic evaluation of method accuracy should address the effects that all of these variables have on the sensitivity and specificity of detecting isotope incorporators. Since DNA-SIP experiments are costly, technically difficult, and laborious, it is not practical to perform empirical assessment across this full range of variables.

Fortunately, the physics of isopycnic centrifugation have been well characterized mathematically, and the behavior of individual DNA fragments in CsCl gradients is highly reproducible and predictable from first principles (Meselson et al., 1957; Fritsch, 1975; Birnie and Rickwood, 1978). In addition, genome sequences are available for thousands of diverse microorganisms, and these genomes can be used to generate DNA fragments representative of community DNA (Youngblut and Buckley 2014). Hence, we can simulate realistic HTS-DNA-SIP data for *in silico* microbial communities that differ in diversity (richness, evenness, and composition), where the relative abundance, genome G + C content, and atom % excess isotope are defined for discrete DNA fragments from every genome. We have developed a computational toolset for simulating HTS-DNA-SIP data (SIPSim) and used this simulation framework to systematically and objectively evaluate how changes in key SIP experimental parameters affect HTS-DNA-SIP accuracy.

## Results

### Model validation and parameter estimation

The SIPSim model starts with a set of user-designated genomes and user-designated experimental parameters (*e.g.* number of gradient fractions, desired community characteristics, desired isotopic labeling characteristics) as described (see Experimental Procedures and *Supporting Information*). Briefly, the genomes are fragmented as would occur during DNA extraction, isotopic labeling is applied to some number of genomes as specified by the user, the BD distributions are determined for each DNA fragment and fragment collections are then binned into gradient fractions, fragments are sampled from each fraction as would occur during amplification and DNA sequencing of SSU rRNA genes, and then the relative abundance is calculated for each OTU (Figure 1). The model produces results that are highly similar to those observed in empirical experiments, including the ability to detect DNA fragments throughout the density gradient (Figure 2).

**Figure 1.**
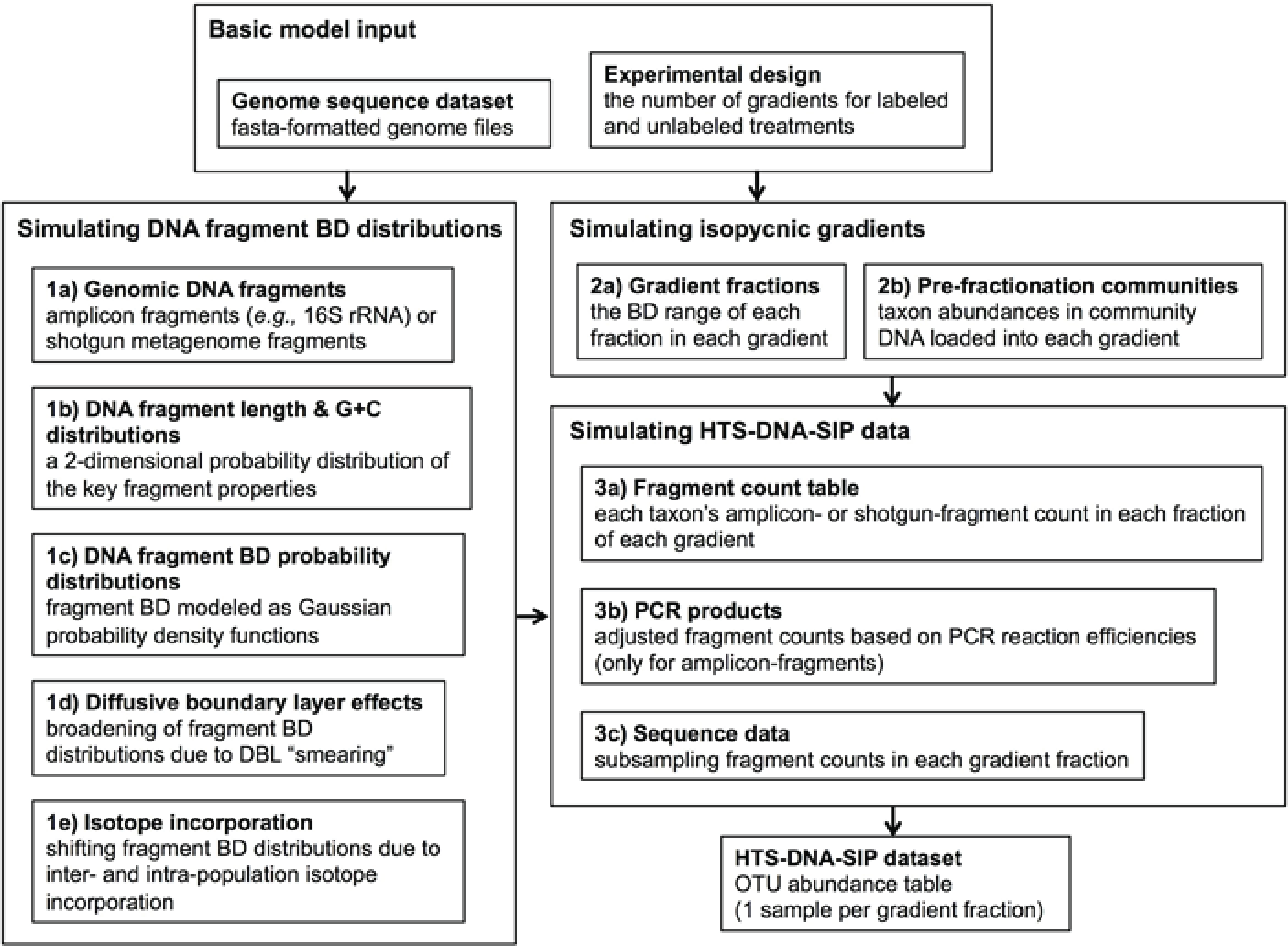
The SIPSim simulation workflow involves three major stages, which are broken down into multiple steps. Stage 1 involves generating a buoyant density distributiony of gDNA fragments for each genome. Stage 2 involves simulating the isopycnic gradients for a particular experimental design. Stage 3 involves generating a HTS-DNA-SIP dataset based on the fragment BD value distributions simulated in Stage 1 along with the isopycnic gradient data generated in Stage 2. The output is a table (“HTS-DNA-SIP dataset”) of taxon relative abundances in each gradient fraction in each gradient. See Experimental Procedures for a more detailed description of the simulation workflow.

**Figure 2.**
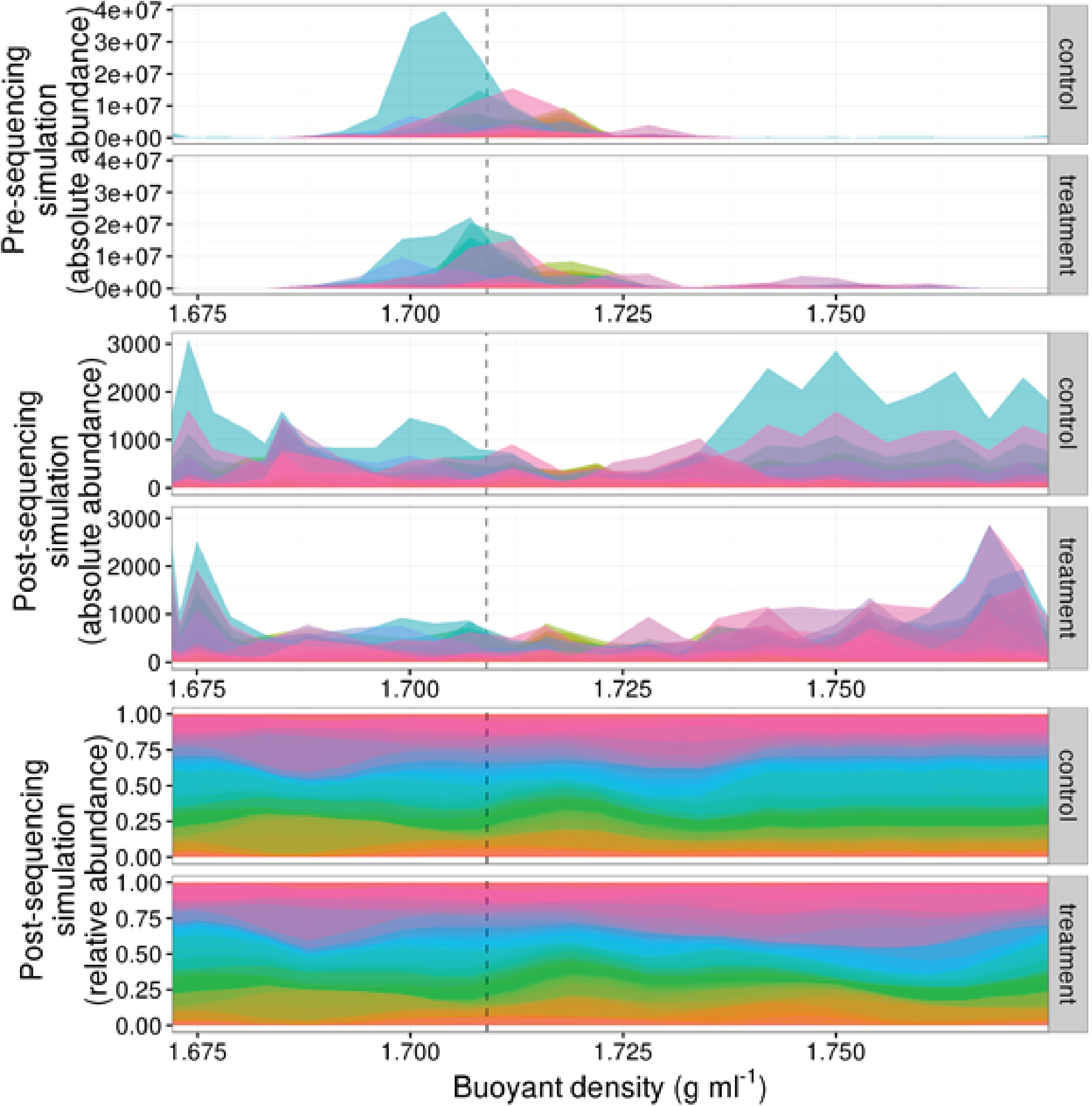
Compositional effects can distort and obscure BD shifts resulting from ^13^C isotope incorporation. The plots show DNA fragment distributions resulting from simulation of 1147 taxa (one color per taxon) within CsCl gradient pairs consisting of: a ^12^C-control (“control”) and a ^13^C-treatment (“treatment”) gradient. For this simulation, all taxa in the control gradient had 0% atom excess ^13^C, while 10% of taxa in the treatment gradient were randomly assigned 100% atom excess ^13^C. “Pre-sequencing simulation” (top) and “Post-sequencing simulation” (middle, and bottom) show fragment BD distributions before and after simulating the effect of uniform random sampling which results from high throughput sequencing of all gradient fractions at an equal number of sequences. The “absolute abundance” (top and middle) indicates the number of DNA fragments from each taxon in each gradient fraction, while “relative abundance” (bottom) indicates the relative abundance of each taxon. Note that the top plot represents the actual amplicon-fragment distributions in an isopycnic gradient at equilibrium, while the bottom plot represents the sampled fragment distributions obtained after high throughput sequencing. The dashed vertical line is provided as a point of reference and designates the theoretical buoyant density of an unlabeled DNA fragment with 50 % G + C (as modeled in Eq. 1)

The development of the simulation model was guided by established centrifugal theory and by comparison of simulated results to empirical data (as in Experimental Procedures in *Supporting Information*). First, we performed a simple evaluation of model performance by recreating results from a prior DNA-SIP experiment with *Methanosarcina barkeri* MS and *Methylobacterium extorquens* AM1 (Lueders et al., 2004) (Figure S1). Simulated DNA distributions (both in terms of total DNA and SSU rRNA gene amplicon copies) significantly and strongly correlated with the empirical data for both taxa (*p* < 0.003 for all comparisons; see Table S1). In addition, the simulated SSU rRNA gene amplicon-fragment BD distributions were shifted 0.007 g ml^−1^ toward the middle of the BD gradient relative to the shotgun-fragments (“total DNA”), a phenomenon also observed in the empirical data. This central tendency for SSU rRNA amplicon-fragments reflects G + C conservation of the *rrn* operon, as previously described (Youngblut and Buckley, 2014).

Next, we determined whether the simulation accurately modeled variation in BD within complex mixtures of unlabeled DNA by comparing simulation results to empirical results obtained with unlabeled DNA from soil. For this purpose we used empirical data from an experiment (Youngblut et al., in prep.) in which DNA was extracted from soil microcosms at 1, 3, 6, 14, 30, and 48 days following the addition of an unlabeled carbon source mixture. These six DNA samples were equilibrated in CsCl gradients, fractionated by BD, and SSU rRNA gene amplicons were sequenced for ~24 fractions from each gradient. Simulation input included 1147 microbial genomes (see Experimental Procedures), hence the soil data was resampled to 1147 OTUs in order to standardize the richness of the simulated and empirical data. Ideally, we could map SSU rRNA sequences from soil to all bacterial genomes available in public databases, but genome composition can vary dramatically across taxa that have identical SSU rRNA gene sequences. Since the genome sequences of taxa in the empirical HTS-DNA-SIP dataset cannot be confidently assigned to genomes in existing databases, a direct mapping of taxa (and their genomes) between the empirical and simulated datasets was not possible. We therefore employed metrics that capture variation in DNA fragment BD distributions within density gradients, and which thereby allow for gradient to gradient comparison of DNA BD distributions (see *Supporting Information*).

The empirical BD distributions (Figure S2) show that temporal change in soil mesocosm community composition caused dramatic shifts in the Shannon diversity of ‘heavy fractions’ even in the absence of isotopic labeling (Figure S2B), with heavy fraction diversity increasing at later time points. Moreover, taxonomic similarity within a gradient is auto-correlated across the BD gradient (Figure S2C). Lastly, variance in amplicon fragment BD is positively correlated with OTU relative abundance in the community (Figure S2D), with highly abundant OTUs found throughout the CsCl gradient. We found that the simulation model was able to recapitulate these results across a wide range of parameter space, and that the variance between simulated and empirical results was less than that observed between replicate empirical samples (Figure S3). We used these comparisons to determine model parameter values (Table S2), which provided the best fit to the actual behavior of DNA fragments in CsCl gradients (as described in *Supporting Information*).

### The influence of isotope incorporation on HTS-DNA-SIP accuracy

We hypothesized that both the number of taxa that incorporate isotope and the atom % isotope incorporation per taxon would substantially affect the accuracy of HTS-DNA-SIP methods. To test these predictions, we simulated HTS-DNA-SIP datasets for both ^13^C-labeled samples and unlabeled controls (3 replicates of each), while varying both the number of incorporators (1, 5, 10, 25, or 50 % of taxa) and the atom % isotope incorporation for each taxon (0, 15, 25, 50, or 100 atom % excess ^13^C). Taxa in the control were always set to 0 % isotope incorporation. Each simulation was replicated 10 times, with differing taxa randomly designated as incorporators in each replicate. We evaluated 4 methods used to analyze HTS-DNA-SIP data: Heavy-SIP, q-SIP, HR-SIP, and MW-HR-SIP. Heavy-SIP involved simply identifying as incorporators all taxa observed in “heavy” gradient fractions of labeled gradients, which provided a baseline of accuracy for the more complex HTS-DNA-SIP analyses. q-SIP and HR-SIP were performed as described in Hungate et al., (2015) and Pepe-Ranney et al., (2016a), respectively. MW-HR-SIP was performed similarly to HR-SIP, but with multiple overlapping “heavy” buoyant density windows (see Experimental Procedures).

As expected, both the number of incorporators and the amount of isotope incorporated affected accuracy (Figure 3). However, the effect of these parameters on specificity and sensitivity varied depending on the analytical method (Figure 3). Specificity is the proportion of true negatives observed out of all true negatives expected, and so specificity declines in direct relation to an increase in the number of false positives. For example, a specificity of 0.8 would generate 200 false positives in a sample of 1000 unlabeled taxa. Specificity, as measured across a wide range in parameters, was highest for MW-HR-SIP (1± 0; ave. ± s.d.) and HR-SIP (1 ± 0), substantially lower for q-SIP (0.88 ± 0.06), and very low for Heavy-SIP (0.28 ± 0.16) (Figure 3).

**Figure 3.**
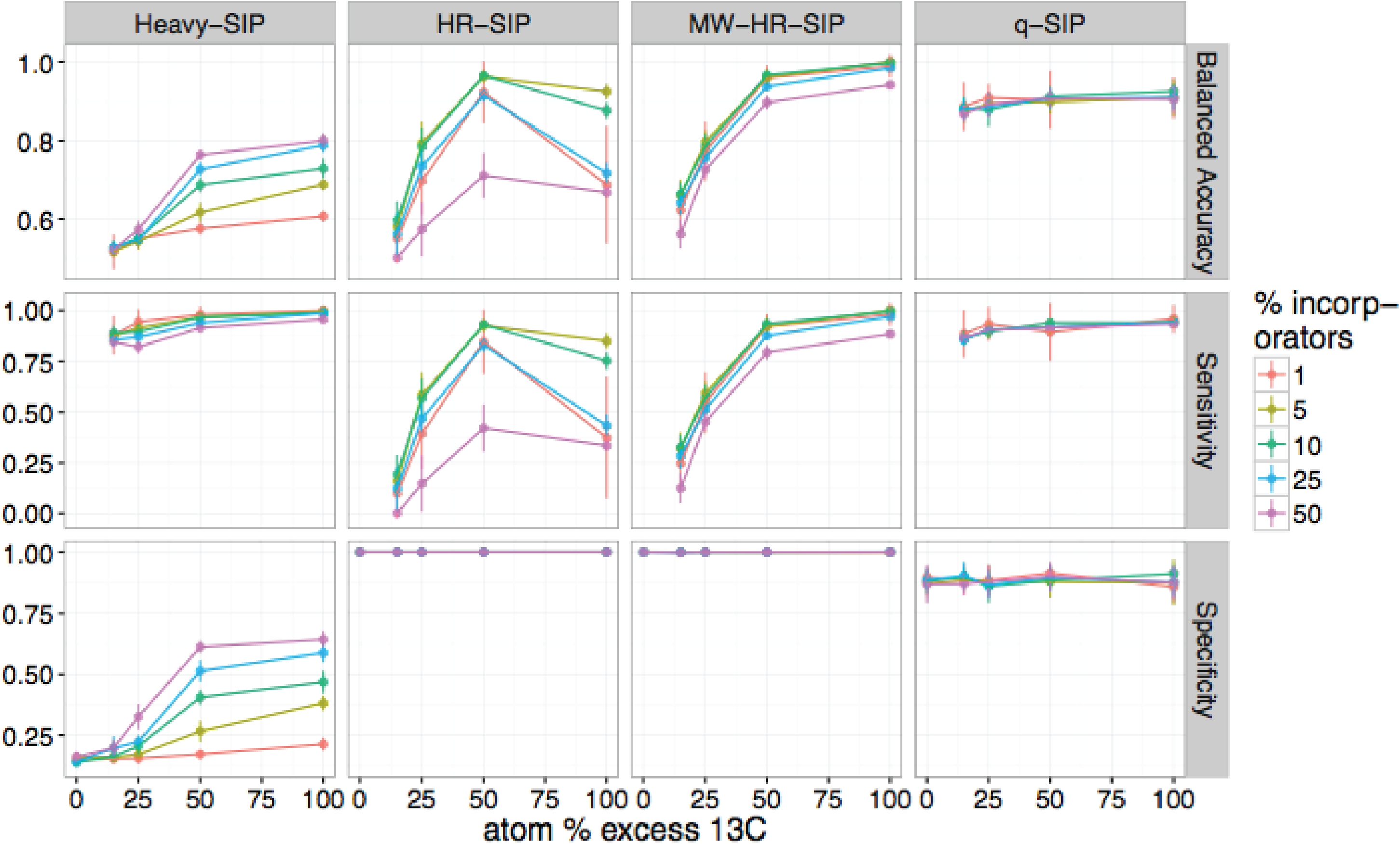
HTS-DNA-SIP methods vary in accuracy depending on the ^13^C atom % excess of DNA and the number of taxa that incorporate isotope. Points and bars represent means and standard deviations, respectively (n = 10 simulations). Specificity indicates the proportion of true negatives that are identified correctly and it is used to quantify false positives. Sensitivity indicates the proportion of labeled taxa (true positives) identified correctly. Balanced accuracy is a function of both specificity and sensitivity. The x-axis indicates the amount of ^13^C isotope present in taxa that are labeled, and different colors are used to indicate the percentage of taxa that have incorporated ^13^C as indicated by the legend.

Sensitivity is the fraction of true positives observed out of all true positives expected. For example, a sensitivity of 0.7 means that a method failed to detect 30 % of the incorporators present. Both q-SIP and Heavy-SIP had relatively high sensitivity (median values of 0.91 and 0.93, respectively), and the sensitivity of these methods was largely insensitive to the atom % excess of DNA and the number of incorporators (Figure 3). In contrast, the sensitivities of both HR-SIP and MW-HR-SIP were highly responsive to the atom % excess of DNA, and the number of incorporators (Figure 3). For these methods, sensitivity declined in proportion to the atom % excess ^13^C label in DNA.

Balanced accuracy is calculated as the mean of specificity and sensitivity. We observed a tradeoff in balanced accuracy in relation to the atom % excess ^13^C of DNA. MW-HR-SIP had the highest accuracy when % atom excess ^13^C exceeded 50 %, but q-SIP had higher accuracy at lower levels of isotope incorporation (Figure 3). This tradeoff in balanced accuracy resulted from a difference in the tolerance for false positives. For example, MW-HR-SIP produced nearly zero false positives but as a result of its high specificity, it lost sensitivity at lower levels of isotope incorporation. In contrast, q-SIP detected labeled taxa across a wider range of isotope incorporation, but it did so at the cost of a large number of false positives.

### The influence of community variation on HTS-DNA-SIP accuracy

All HTS-DNA-SIP analyses rely upon comparisons made between isotopically enriched experimental treatments and their corresponding unlabeled controls. In real SIP experiments the composition of replicate post incubation communities are likely to vary somewhat as a result of sample heterogeneity and incubation effects. However, the simulations described above assume random sampling from identical pre-fractionation (post-incubation) community structures. We hypothesized that an increase in variation in community composition between treatment and control samples would decrease the accuracy of HTS-DNA-SIP analyses. To test this hypothesis, we generated simulations in which isotope incorporation was held constant (100 atom % excess ^13^C; 10 % of OTUs are incorporators) but beta-diversity was varied among 3 replicate treatment and 3 replicate control samples. We varied beta-diversity in two ways: *i*) using permutation to vary the rank abundance of a fixed proportion of community members and *ii*) varying the proportion of taxa shared between communities. For each simulation scenario, we calculated the mean Bray-Curtis distance among communities in order to provide a real-world metric for gauging the potential accuracy of actual DNA-SIP experiments.

As hypothesized, increased beta-diversity among samples had a substantial impact on the accuracy of HTS-DNA-SIP methods (Figure 4). Accuracy was impacted more by the number of taxa shared between samples than by differences in taxon abundance (Figure S4). The sensitivity of q-SIP declined as beta-diversity increased, falling to 0.64 ± 0.04 (ave. ± s.d.) when samples shared 80 % of their OTUs (Figure S4). In contrast, the sensitivities of MW-HR-SIP and Heavy-SIP were least affected by changes in beta-diversity and these methods had the highest sensitivity overall (0.81 ± 0.04 and 0.82 ± 0.03 at 80 % shared OTUs, respectively; ave. ± s.d.). Increasing the beta-diversity of samples had little effect on the specificity of q-SIP but diminished slightly the specificity of HR-SIP and MW-HR-SIP (Figure 4). Despite these declines, HR-SIP and MW-HR-SIP maintained specificity that was greater than or equal q-SIP and Heavy-SIP throughout most parameter space (Figure 4).

**Figure 4.**
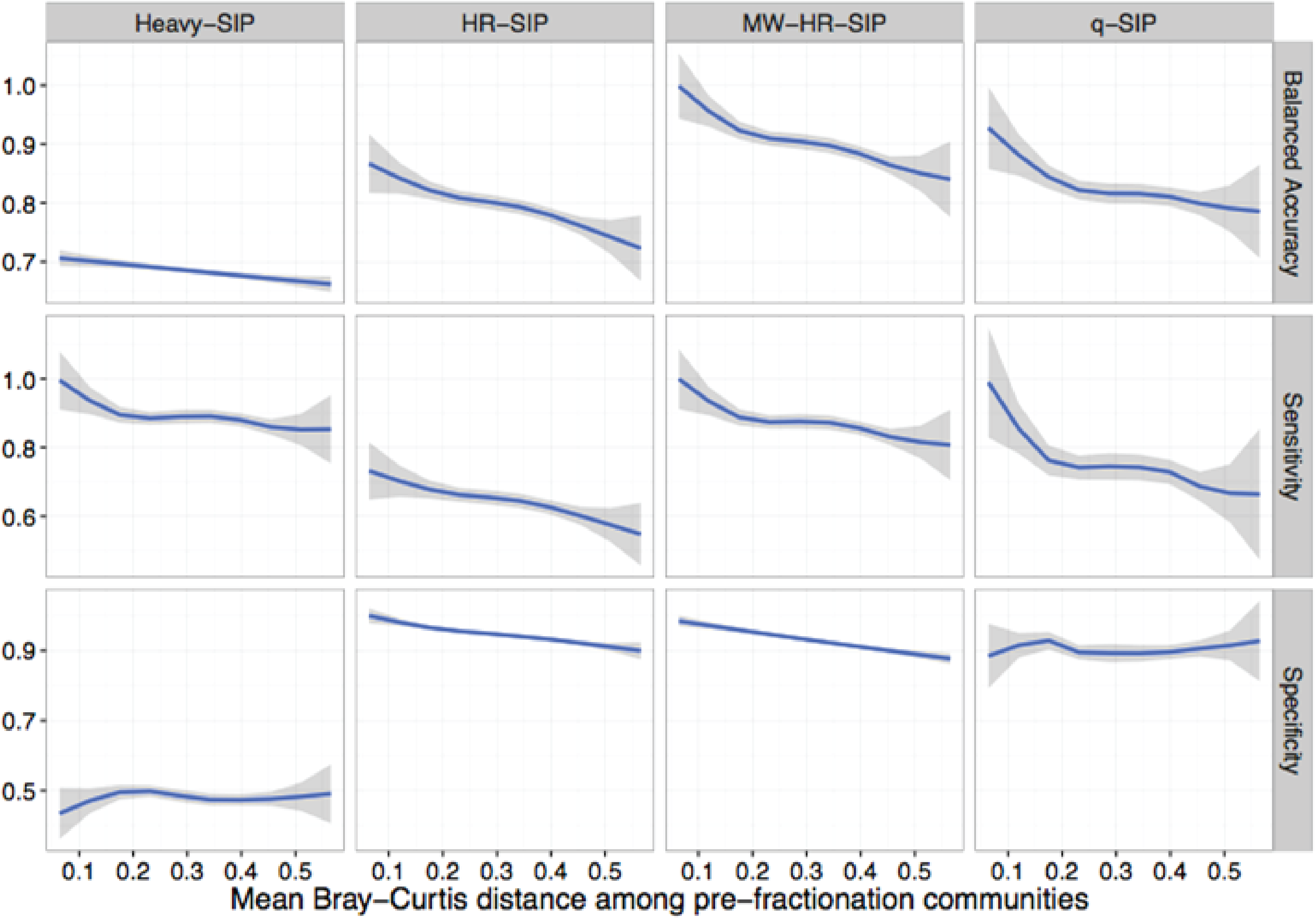
HTS-DNA-SIP methods differ in their sensitivity to community dissimilarity between replicate samples. Beta diversity, expressed as Bray-Curtis dissimilarity, was varied between simulated replicates (3 replicates each for ^12^C-control and ^13^C-treatment gradients) to determine the effect that community dissimilarity between replicates has on method accuracy. Variation in beta diversity was simulated by systematically varying two parameters: the percent of taxa shared between replicate samples (80, 85, 90, 95, or 100%) and the percent of taxa whose rank abundances that were permuted (0, 5, 10, 15, or 20%), with 10 simulation replicates for each parameter set. The blue lines are LOESS curves fit to accuracy values for all simulations (*n* = 250), and the grey regions represent 99% confidence intervals. For all simulations, 10% of the community were incorporators (100% atom excess ^13^C).

Of the methods evaluated, MW-HR-SIP had the highest balanced accuracy across the widest range of parameters tested (Figure 4). Regardless, the balanced accuracy for MW-HR-SIP was negatively affected by an increase in beta-diversity, falling from 0.98 ± 0.02 to 0.86 ± 0.02 (ave. ± s.d.) when the Bray-Curtis dissimilarity between samples was increased beyond 0.5 (Figure 4). These results highlight the overall negative impact that sample-to-sample variation has on HTS-DNA-SIP accuracy, and the importance of minimizing experimental variation between unlabeled controls and labeled treatments in SIP experiments.

### Using HTS-DNA-SIP data to quantify atom % excess

So far we have focused on the accuracy of HTS-DNA-SIP methods with respect to the identification of taxa that incorporate isotope into their DNA. However, changes in DNA BD can also be used to quantify the isotope enrichment of DNA from particular taxa. Two approaches have been used to evaluate isotope enrichment from HTS-SIP data: q-SIP and ΔBD, with the latter being a complementary analysis to HR-SIP (Pepe-Ranney et al., 2016a). Both ΔBD and q-SIP derive quantitative estimates from measuring taxon BD shifts (and thus atom % excess) in the labeled treatment gradient(s) versus their unlabeled counterparts. The ΔBD method attempts to measure the extent of the BD shift directly from the compositional sequence data, while q-SIP utilizes relative abundances transformed by qPCR counts of total SSU rRNA copies. Therefore, ΔBD accuracy likely suffers from compositional effects inherent to HTS datasets, while q-SIP accuracy is dependent on qPCR accuracy and variation.

We assessed the quantification accuracy of both methods using the simulations described previously, where either the amount of isotope incorporation or sample beta-diversity was varied. We found that ΔBD produced estimates of isotope incorporation that were closer on average to the true value compared to q-SIP, but ΔBD values had much higher variance than q-SIP estimates (Figure 5). Furthermore, the variance in ΔBD atom % excess ^13^C estimates increased substantially with even moderate increases in beta-diversity between samples, while the q-SIP estimations were largely invariant across the simulation parameter space (Figure S5A). However, mean q-SIP values consistently underestimated the true ^13^C atom % excess by 30.2-39.2% (Figure 5B). Overall, quantitative estimates of isotope incorporation for individual taxa were less variable with q-SIP, though q-SIP consistently miss-estimated actual levels of isotope enrichment.

**Figure 5.**
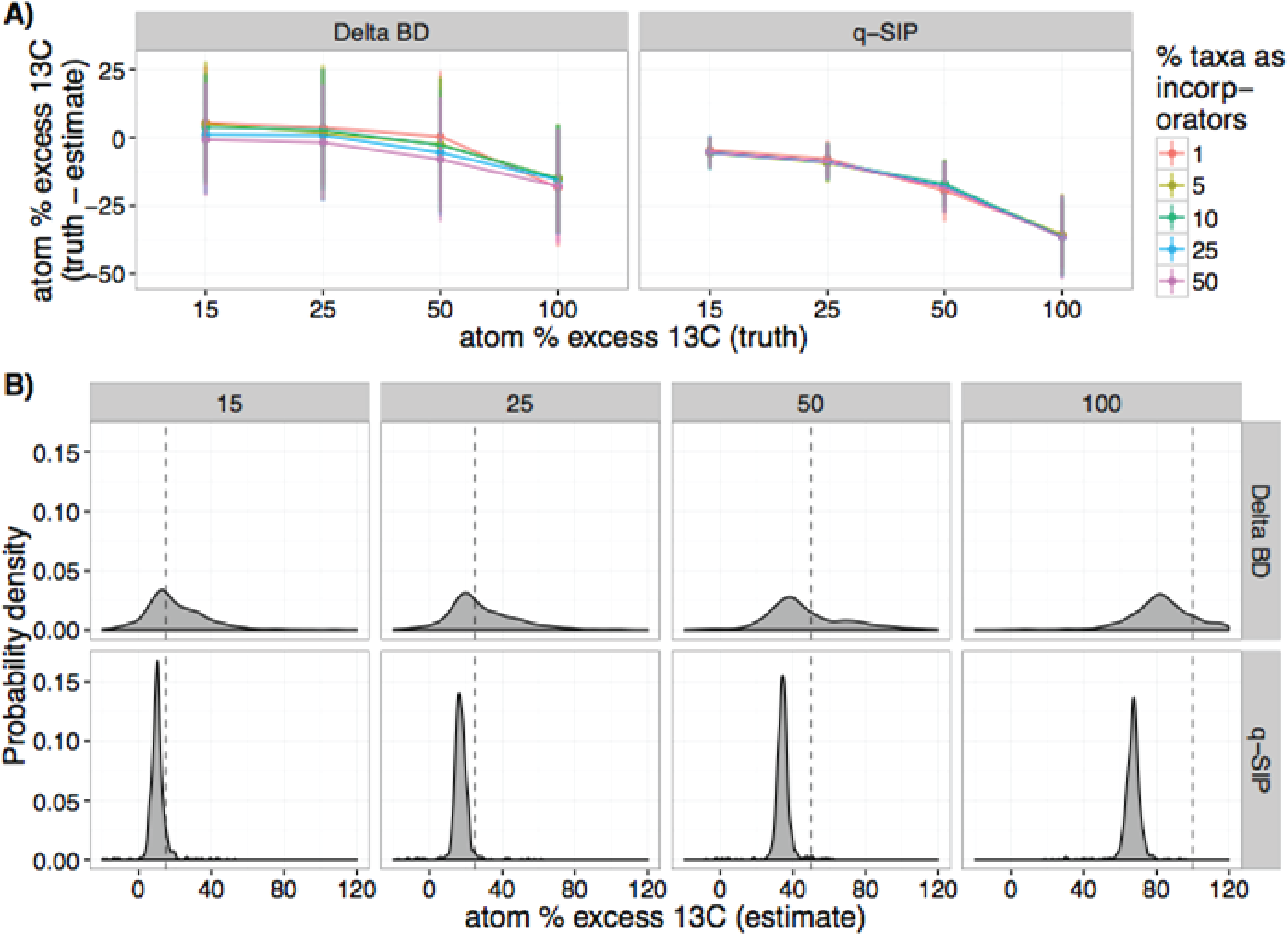
ΔBD and qSIP vary in their accuracy at estimating ^13^C atom % excess of labeled DNA fragments. (A) The accuracy of both methods declines as the amount of ^13^C in DNA increases, but accuracy is not affected by the percent of taxa that are labeled; values indicate the mean and standard deviation (n = 10 simulations). (B) Probability density plots indicate that estimates of ^13^C atom % excess made using *ΔBD* have greater variance than those made using *qSIP*, but both estimates systematically underestimate levels of isotope incorporation. Each vertical pair of panels indicates the probability density for estimates made across different levels of isotope incorporation (15, 25, 50, and 100 atom % excess), and the dashed line indicates the actual level of isotopic enrichment. For the calculation of probability density, 10 % of taxa were labeled using the level of enrichment indicated in each panel.

## Discussion

Our simulation framework (SIPSim) provides a tractable platform for evaluating the accuracy of DNA-SIP methods and for developing new methods to analyze DNA-SIP data. Given the laborious nature of DNA-SIP experiments, it is impossible to use empirical analyses with mock communities to evaluate the range of parameter values that can be investigated through simulation. In addition, both the physics of density gradient centrifugation and the physical properties of genomic DNA are well established, making the simulation of DNA-SIP data both tractable and reliable. Without rigorous assessment of DNA-SIP methods, it is difficult to determine the likelihood of false negatives (Type II error) and false positives (Type I error) across the wide range of experimental conditions in which DNA-SIP has been employed in the literature. Issues of Type I and Type II statistical error are compounded in the analysis of HTS-DNA-SIP by the nature of HTS data, where it is necessary to make many thousands of comparisons to identify those OTUs that change in response to treatment. This multiple comparison problem has major implications for statistical power and the likelihood of false detection (Paulson et al., 2013). We have used SIPSim to test the effects of multiple parameters on the accuracy of current methods for analyzing HTS-DNA-SIP data. Furthermore we used observations from the model to develop MW-HR-SIP, an analytical approach with balanced accuracy higher than any other current method, with a higher sensitivity than HR-SIP, a higher specificity than q-SIP and Heavy-SIP, and a higher robustness to inter-sample beta-diversity than all other currently available methods

Although both HR-SIP and q-SIP use high throughput SSU rRNA amplicon sequencing of many gradient fractions, their different approaches for detecting isotope incorporators result in substantial differences in sensitivity and specificity. In q-SIP, taxon relative abundance is transformed using qPCR data to estimate counts of SSU rRNA gene copies across gradient fractions; however, this approach resulted in a large number of false positives (8 ± 0.3 to 15 ± 0.7 % of the unlabeled taxa evaluated were misidentified as labeled; Figure 3 and Figure 4). Moreover, the number of false positives detected by q-SIP increased dramatically in response to variation in community structure between samples (Figure 4). In contrast, HR-SIP had negligible false positives under a wide range of parameters (Figure 3 and Figure 4), but had lower sensitivity (more false negatives) than other methods. However, we found that the sensitivity of HR-SIP was improved without compromising specificity by using a multi-window analysis (MW-HR-SIP) in place of the single window analysis used in HR-SIP. MW-HR-SIP has high sensitivity and specificity across a range of experimental parameters, provided that the atom % excess ^13^C of DNA is in the range of 50-100% (Figure 3). At lower levels of isotope incorporation, q-SIP has better sensitivity, but this sensitivity comes at the cost of detecting a large number of false positives (*i.e.* low specificity). This tradeoff between specificity and sensitivity can be contextualized by considering a community that contains 1045 unlabeled taxa and 55 labeled taxa. If these 55 taxa are labeled at 50% atom excess ^13^C, both methods do a good job detecting truly labeled taxa (MW-HR-SIP: 51 ± 2; q-SIP: 50 ± 2), but q-SIP detects far more false positives (MW-HR-SIP: 0 ± 1; q-SIP: 126 ± 8). If these 55 taxa are instead labeled at 25% atom excess ^13^C then MW-HR-SIP detects fewer labeled taxa than q-SIP (MW-HR-SIP: 33 ± 3; q-SIP: 50 ± 2), but q-SIP still detects far more falsely labeled taxa (MW-HR-SIP:1 ± 0; q-SIP: 122 ± 8). In these examples, about 71% of the taxa identified by q-SIP as labeled are actually unlabeled.

When considering the relative importance of sensitivity versus specificity for DNA-SIP experiments, the ability to detect taxa that incorporate isotope is only useful if those identifications can be made with high confidence (*i.e*. with a low number of false positives). Therefore, based on our results, MW-HR-SIP is the most robust method for identifying isotope incorporators from HTS-DNA-SIP data. In addition to its high specificity and better ability to handle variance between replicate samples, MW-HR-SIP has the added advantage of not requiring qPCR to be performed on each gradient fraction. It should be noted, however, that the primary objective for which MW-HR-SIP was designed is the accurate detection of labeled taxa, regardless of level of isotopic enrichment, while a major goal of q-SIP is to quantify the atom % excess of individual taxa.

In regards to methods used to quantify the atom % excess of individual taxa from HTS-DNA-SIP data, we found that the utility of q-SIP or ΔBD varied depending on the hypothesis being evaluated. ΔBD produced more accurate estimates of mean ^13^C atom % excess than q-SIP (Figure 5 and Figure S5), and so this approach may be suitable when seeking to make relative comparisons in the degree of labeling between large groups of taxa (as described in Pepe-Ranney, et al., 2016). However, the high variability of this approach causes ΔBD to be unreliable in determining differences in atom % excess ^13^C at the scale of individual OTUs. Alternatively, q-SIP produced much more stable estimates of atom % excess ^13^C among individual taxa, but the method resulted in systematic underestimates of isotope incorporation

The SIPSim framework makes it possible to both evaluate hypothetical outcomes of DNA-SIP experiments before they are performed and to evaluate the accuracy of HTS-DNA-SIP data analysis methods. For brevity, we have only focused on a few key variables that could affect the accuracy of HTS-DNA-SIP methods. However, SIPSim can also be used to assess the accuracy of DNA-SIP methods across a range of possible real-world scenarios. For instance, spatial or population-level heterogeneity could result in taxa that are not homogeneously labeled (Lennon and Jones, 2011). Such systematic heterogeneity in labeling would manifest as “split” (bimodal or multimodal) distributions of DNA fragments in an isopycnic gradient. It would be challenging to evaluate such scenarios empirically, but SIPSim can be readily used to evaluate a range of such scenarios. SIPSim can also be used to evaluate the effect of sequencing depth on the statistical power needed to resolve isotope incorporation in rare taxa. Such information should be useful in planning HTS-DNA-SIP experiments, to ensure that the experiment has a reasonable chance of success before it is performed. Finally, SIPSim provides a toolkit for developing and improving analytical methods used in DNA-SIP experiments. For example, a hybrid method that combines aspects of MW-HR-SIP and q-SIP may be able to produce robust incorporator identification while also providing accurate estimates of the atom % excess of individual taxa.

## Conclusion

With our newly developed simulation toolset, we determined that MW-HR-SIP has the highest accuracy of currently available methods for identifying taxa that have incorporated isotope in HTS-DNA-SIP experiments. The use of MW-HR-SIP resulted in a negligible number of false positives and its ability to detect true positives varied in relation to the isotopic enrichment of DNA. Generally, we found that the specificity of all HTS-DNA-SIP methods declined with increased beta-diversity among replicate samples. Thus, given that accuracy declined most rapidly between a mean Bray-Curtis distance of 0 and 0.2 for all methods evaluated (Figure 4), we recommend that researchers strive for mean Bray-Curtis distances of <0.2 among replicate samples used in SIP experiments (*i.e.* between treatments and their corresponding controls).

## Experimental Procedures

### Theory underlying the simulation framework

DNA stable isotope probing employs isopycnic centrifugation to separate isotopically enriched (“heavy”) DNA molecules from unlabeled (“light”) DNA based on their differences in buoyant density (BD). Isopycnic centrifugation is distinguished from other centrifugation methods in that centrifugation is carried out long enough to both generate a density gradient (typically using CsCl for DNA-SIP) and have all macromolecules of interest reach sedimentation equilibrium, which is the point at which sedimentation rates equal rates of diffusion (Hearst and Schmid, 1973; Birnie and Rickwood, 1978). Empirical studies have shown that the average BD (ρ) of a mixture of DNA molecules is linearly related to the average G + C content for that collection of molecules:

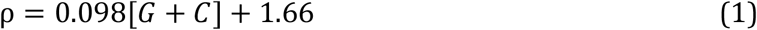

where [G + C] is the mole fraction of G+C content (Schildkraut et al., 1962; Birnie and Rickwood, 1978). In addition, empirical studies have also shown that homogeneous mixtures of DNA molecules form a Gaussian distribution in an isopycnic gradient when at sedimentation equilibrium (Meselson et al., 1957; Fritsch, 1975). Therefore, in order to model the BD distribution of a heterogeneous set of genomic DNA fragments, a Gaussian distribution must be estimated for each homogeneous subset of molecules rather than using discrete BD values (as described in *Supporting Information*). Based on the work of Meselson and colleagues (Meselson *et al.*, 1957), Fritsch (1975) derived an equation describing time to reach sedimentation equilibrium, which can be reworked to calculate the standard deviation (σ) of the Gaussian distribution:

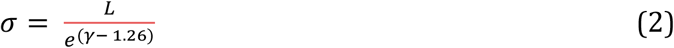

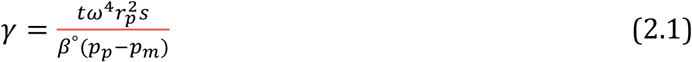

where *L* is the effective length of the gradient (cm), *t* is time in seconds, *ω* is the angular velocity (radians sec^−1^), *r_p_* is the distance of the particle from the axis of rotation (cm), *s* is the sedimentation coefficient of the particle, *β*° is the coefficient specific to the density gradient medium (*e.g.* CsCl); *p_p_* and *p_m_* are the maximum and minimum distances between the gradient and axis of rotation (cm) (Fritsch, 1975). By assuming that sedimentation equilibrium has been reached for all macromolecules of interest, Clay and colleagues derived a simplified equation for determining σ from the calculations in (Schmid and Hearst, 1972):

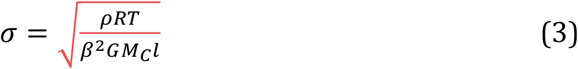

where *ρ* is the BD of the particle, *R* is the universal gas constant, *T* is the temperature in Kelvins, *β* is a proportionality constant for aqueous salts of specific densities, *G* is a buoyancy factor as described in (Clay et al., 2003), *M_C_* is the molecular weight per base pair of DNA, and *l* is the fragment length (bp). For most DNA-SIP experiments, the assumption of sedimentation equilibrium for all DNA fragments is likely to be unrealistic for relatively short DNA fragments (*e.g*. < 4 kb), given that the time to equilibrium rises dramatically with decreasing fragment length (Meselson et al., 1957; Birnie and Rickwood, 1978; Youngblut and Buckley, 2014). However, the ultracentrifugation durations used in typical DNA-SIP experiments should still generally produce small σ values for short DNA fragments according to Eq. 2 (Neufeld et al., 2007). Therefore, equation Eq. 3 provides a good approximation for modeling the BD distribution of DNA in density gradients generated in typical DNA-SIP experiments.

The distribution of a heterogeneous mixture of DNA fragments in an isopycnic gradient can thus be modeled by integrating the Gaussian distributions of each homogeneous subset of DNA fragments, where the mean of each Gaussian is determined by Eq. 1 and the standard deviation derived from Eq. 3. In this way, the BD distribution for a given genome in an isopycnic gradient can be modeled by the following steps: simulate genome fragmentation resulting from DNA extraction, bin gDNA fragments with respect to length and G + C content, model Gaussian distribution for each fragment bin, and then integrate these distributions to describe the cumulative DNA distribution in the gradient.

We found that empirical DNA fragment distributions differed from the expectations of a strictly Gaussian model (Figure S2), and we determined that these differences could be reconciled on the basis of established principles of fluid mechanics (as described below and in *Supporting Information*). Based on empirical measurements, we found that most taxa with relative abundances > 0.1% are detected in all gradient fractions when unlabeled DNA is subjected to CsCl gradient centrifugation and SSU rRNA amplicon sequencing is performed across a wide range of density gradient fractions (Figure S2). This observation is in general congruence with observations in the literature (Birnie and Rickwood, 1978; Lueders et al., 2004; Leigh et al., 2007), but it does not match the expectation that DNA fragment distributions are strictly Gaussian, since the Gaussian model predicts that DNA fragments should be undetectable (i.e. probability density < 1e^−7^) at either end of the density gradient (Figure S6). We explain the difference between observed and expected DNA distributions as a function of fluid mechanics during gradient reorientation.

During isopycnic centrifugation, the buoyant density gradient forms perpendicular to the axis of rotation (Figure S7), and gradient reorientation during centrifuge deceleration is dramatic, especially for vertical rotors (Flamm et al., 1966). While the distortion of the BD gradient during reorientation has been shown to be minimal in the aggregate (Fisher et al.,1964; Flamm et al., 1966), the inevitable presence of a diffusive boundary layer along the tube wall is sufficient to entrain quantities of DNA, which are small but should be readily detectable by high throughput sequencing methods. The flow field that occurs during gradient reorientation entrains along the tube wall a volume with a dimension proportional to flow velocity, fluid viscosity, and surface topography (Tritton, 1977; Cohen and Dowling, 2012). Following gradient reorientation, DNA from the entrained volume will combine with DNA from the reoriented volume, thereby introducing a small amount of non-BD-equilibrium DNA into each gradient fraction (Figure S7). The ability of the diffusive boundary to introduce non-BD-equilibrium DNA into gradient fractions can be modeled as a function of rotor geometry (Figure S7). Assuming sedimentation equilibrium, BD (ρ) can be directly related to the distance from axis of rotation (Birnie and Rickwood, 1978):

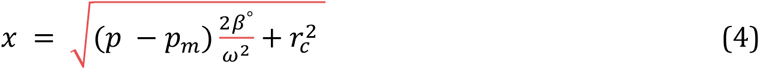

From this calculation, the location of DNA molecules in the centrifuge tube, both during centrifugation and fractionation, can be ascertained by using simple trigonometry along with knowledge of centrifuge tube dimensions and angle to the axis of rotation. A full description of the calculations along with an example can be found at https://github.com/nick-youngblut/SIPSim. The fraction of a taxon's DNA fragments that are in the boundary layer (*D_ti_*) is modeled as:

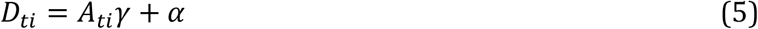

where *A_ti_* is the pre-fractionation community relative abundance of taxon *t* in gradient *i*, γ is a weight parameter determining the contribution of *A_ti_* to *A_b_*, and *α* is the baseline fraction DNA in *A_b_*.

Assimilation of the commonly used isotopes ^13^C and ^15^N into genomic DNA produces linear shifts in BD, with a maximum shift of 0.036 and 0.016 g ml^−1^, respectively (Birnie and Rickwood, 1978). Thus the shift in BD (ρ) can be modeled as:

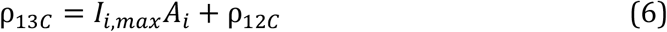

where *I_i,max_* is the maximum possible BD shift if 100% atom excess for isotope *i*, *A* is the atom % excess of isotope *i*, and *ρ12C* is the buoyant density at 0% atom excess.

### SIP data simulation framework overview

Based on the theory described above, our SIP data simulation framework simulates the distribution of gDNA fragments in isopycnic gradients at sedimentation equilibrium. Furthermore, it generates the HTS-DNA-SIP datasets obtained from fractionating isopycnic gradient(s) and performing high throughput sequencing on many of the gradient fractions. Our framework also implements all of the HTS-DNA-SIP analysis methods assessed in this study (Heavy-SIP, HR-SIP, MW-HR-SIP, q-SIP, and ΔBD) and evaluates their accuracy of identifying incorporators or quantifying BD shifts. An overview of our simulation framework is shown in Figure 1.

Our simulation framework is a modular collection of steps that can be grouped in workflow stages that are further broken down into steps (Figure 1). The input is a set of reference genomes in fasta format and a text file designating the experimental design, which includes the number of gradients for labeled treatments and unlabeled controls

Stage 1 involves generating a BD distribution of gDNA fragments for each genome. Step 1a involves simulating the pool of gDNA fragments that is extracted from SIP incubation samples and then loaded into the isopycnic gradients. If amplicon sequence data (*e.g*. SSU rRNA) is to be generated, amplicons from only the fragments containing the PCR template (“amplicon-fragments”) are sequenced, while shotgun metagenomic sequencing can target all gDNA fragments (“shotgun-fragments”). If ≥ 1 PCR primer set is provided, amplicon-fragments are generated from genomic regions fully encompassing genome locations that produced amplicons by *in silico* PCR. Alternatively, shotgun-fragments are randomly generated from all possible genomic locations. The fragment size distribution is user-defined (Table S2)

As described in Eq. 1 & 3, the length and G + C content of a DNA fragment can be used to calculate a probability distribution of its location in the gradient, assuming sedimentation equilibrium. Step 1b uses the fragments simulated in Step 1a to generate a 2-dimensional Gaussian kernel density estimation (KDE) for each taxon, which describes the joint probability of obtaining fragments with a certain length and G + C content from that taxon. From this 2D-KDE, a large number of [length, G + C] vectors can be simulated efficiently for more precise estimations of the fragment BD distributions. Fragment BD distributions are calculated for each taxon in Step 1c by sampling [length, G + C] vectors from the 2D-KDE and calculating Gaussian distribution from each, where the mean is based on Eq. 1 and the standard deviation based on Eq. 3. The collection of Gaussian distributions for all fragments for each taxon is integrated into a BD distribution for all fragments of a taxon with Monte Carlo error estimation, which involves sampling BD values from the collection of Gaussian distributions and estimating a probability density function (PDF) of the fragment BD distribution as a one-dimensional Gaussian KDE. The result is a list of KDEs, with each describing the probability of detecting the gDNA fragments of a taxon at any point along the isopycnic gradient. These fragment BD distributions are modified in steps 1d and 1e by adding diffusive boundary layer (DBL) effects (see Theory) and isotope incorporation, respectively. The “smearing” due to DBL effects is modeled as a uniform distribution describing the increased fragment BD uncertainty, and this uncertainty is integrated into the fragment BD distributions by Monte Carlo error estimation as in Step 3b. The BD shift due to isotope incorporation is modeled in a similar manner, except BD uncertainty is a result of inter- and intra-population variation in the amount of isotope incorporated. Variation of isotope incorporation is modeled as a hierarchical set of mixture models (weighted sets of standard distributions; such as two Gaussians), where the parameters for intra-population mixture models that describe the amount of isotope incorporated by each individual are themselves defined by inter-population mixture models that describe how isotope incorporation varies among taxa.

Stage 2 involves simulating the isopycnic gradients for a particular experimental design. Step 2a involves simulating the BD range size of each fraction of each gradient. Sizes are drawn from a user-defined distribution. Step 2b involves simulating the relative abundance distribution of taxa in the gDNA pools loaded into each gradient (“pre-fractionation communities”). The abundance distribution of each pre-fractionation community is user defined and can vary among gradients. Furthermore, the amount of taxa shared or rank-abundances permutated among communities (*i.e*. the beta-diversity) is user-defined

Stage 3 involves generating a HTS-DNA-SIP dataset based on the fragment BD distributions simulated in Stage 1 along with the isopycnic gradient data generated in Stage 2. In Step 3a, an OTU (taxon) abundance table is generated by sampling from the fragment BD distributions of each taxon generated in Stage 1, with sampling depth determined by pre-fractionation community abundances simulated in Step 2b. The subsampled fragments are then binned into gradient fractions simulated in Step 2a. The resulting OTU table lists the number of gDNA fragments of each taxon in each gradient fraction in each gradient. If the simulated fragments are amplicons, then PCR amplification efficiency biases are simulated in Step 3b based on the PCR kinetic model described in Suzuki and Giovannoni (1996). The model assumes that efficiencies decrease as the product concentration increases due to an increased propensity of single stranded products to re-anneal to their homologous complements. Sequence data is simulated in Step 3c by subsampling from the table of fragment counts (the DNA fragment pool), which produces a final table (“HTS-DNA-SIP dataset”) of taxon relative abundances in each gradient fraction in each gradient.

### SIP data simulation framework parameters

Unless stated otherwise, we made the following assumptions for all simulations in this study. Community abundance distributions were simulated as lognormal distributions with a mean of 10 and a standard deviation of 2. All taxa were shared among communities, and no rank-abundances were permuted (unless otherwise stated as for when evaluating beta-diversity effects). The total number of fragments in each gradient was 1e^9^. Gradient fragment BD range sizes were sampled from a normal distribution, with a mean of 0.004 and a standard deviation of 0.0015. SSU rRNA amplicon-fragments were simulated using the V4-targeting 16S rRNA primers: 515F and 927R (5′-GTGYCAGCMGCMGCGGTRA-3′; 5′-CCGYC AATTYMTTTRAGTTT-3′), as used by Pepe-Ranney and colleagues (Pepe-Ranney, et al., 2016a). The amplicon-fragment size distribution was a left-skewed normal distribution with a mean of ~12 kb, which is similar to size distributions produced from common bead beating cell lysis methods (Kauffmann et al., 2004; Roh et al., 2006; Thakuria et al., 2008). A total of 1e^4^ amplicon-fragments were simulated per genome, which equated to > 100X coverage for the genomic region of interest. Monte Carlo error estimation was conducted with 1e^5^ sampling replicates. Ultracentrifugation conditions were set as in Pepe-Ranney and colleagues (Pepe-Ranney et al., 2016a), with a Beckman TLA-110 rotor spun at 5.5e^4^ rpm for 66 hours at 20°C and an average density gradient 1.7 g ml^−1^. Inter-population variation in isotope incorporation was binary (either 0 % or X % atom excess), and intra-population variation was set to zero. Two key parameters were estimated from empirical HTS-DNA-SIP data: the bandwidth (smoothing factor) for kernel density estimation, and the gamma parameter in Eq. 5. See Table S2 for a full listing of simulation parameters.

### Implementing HTS-DNA-SIP analyses

The HR-SIP method was performed as described in (Pepe-Ranney et al., 2016a; Pepe-Ranney et al., 2016b). Briefly, we used a “heavy” BD window of 1.71-1.75 g ml^−1^, a sparsity cutoff of 0.25 (*i.e.* OTUs must be present in >25% of samples), a log_2_ fold change null threshold of 0.25, and a false discovery rate cutoff of 10 %. ΔBD was determined as described by Pepe-Ranney and colleagues (Pepe-Ranney et al., 2016a), with OTU abundances linearly interpolated across 20 evenly spaced values across the gradient BD range.

We hypothesized that HR-SIP sensitivity could be improved by altering the “heavy” BD window (1.71-1.75 g ml^−1^) in which sequence composition is compared between treatment and control. We evaluated different approaches and found that the analysis of multiple windows (hereby called “MW-HR-SIP”) resulted in a significant improvement in sensitivity relative to HR-SIP. MW-HR-SIP evaluated sequence composition within BD windows of: 1.70-1.73, 1.72-1.75, 1.74-1.77 g ml^−1^ (Figure S8) while adjusting for multiple comparisons q-SIP was conducted as described in Hungate and colleagues (Hungate et al., 2015), with 90 % confidence intervals calculated from 1000 bootstrap replicates. The variance among qPCR replicates was modeled based on the qPCR data provided in Table S2 Hungate et al., (2015). Specifically, we found the qPCR count variance (*σ^2^*) to increase as a function of the mean (*μ*). The following polynomial regression was found to best describe this relationship and was used for simulating all qPCR count values:

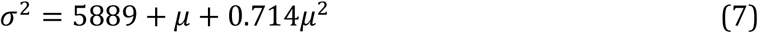

where *μ* was set as the total number of simulated DNA fragments in the gradient fraction(designated in the OTU table from Step 4a)

### Datasets

The genome dataset used to simulate genomic DNA fragments was obtained from Genbank (Benson et al., 2008). From a list of all bacterial genomes designated as “complete”, one representative was chosen per species in order to reduce the bias toward highly represented species. We found the dataset to contain a rather high proportion (~12 %) of low G + C organisms (< 30 % G + C); most of which were obligate endosymbionts. We randomly sampled a subset of these low G + C genomes in order to reduce the proportion of low G + C organisms to just 1 % of the genome dataset. The resulting dataset consisted of 1147 bacterial genomes.

In order to simulate empirical data from Lueders and colleagues (Lueders et al., 2004), the genome sequences of *Methanosarcina barkeri* MS and *Methylobacterium extorquens* AM1 were downloaded from Genbank. Amplicon-fragments were simulated with the primers Ar109f (5′-ACKGCTCAGTAACACGT-3′), Ar915r (5′-GTGCTCCCCCGCCAATTCCT-3′), Ba519f (5′-CAGCMGCCGCGGTAANWC-3′), and Ba907r (5′-CCGTCAATTCMTTTRAGTT-3′). Atom % excess was assumed to be 100 %, and isopycnic centrifugation conditions were simulated as specified in Lueders et al., (2004).

For model evaluation (see *Supporting Information - Results*), we downloaded the genomes *Clostridium ljungdahlii* DSM 13528, *Escherichia coli* 1303, and *Streptomyces pratensis* ATCC 33331 from Genbank.

The HTS-DNA-SIP dataset from Youngblut and colleagues consisted of SSU rRNA MiSeq sequences (V4 region) of ~24 fractions per gradient from 6 gradients of unlabeled controls (Youngblut et al., in prep.). These data were subsampled to obtain a total richness equal to the 1147 OTUs in our reference genome dataset. The sequence data is available from the NCBI under BioProject PRJNA382302.

### Software implementation

The SIP simulation framework was mostly written in Python v2.7.11, with some accompanying code written in C++ v4.9.2 and R v3.2.3 (R Core Team, 2016). MFEprimer v2.0 was used to perform *in silico* PCR (Qu *et al.*, 2009). The software, along with documentation and examples, can be found at https://github.com/nick-youngblut/SIPSim. All genomes were downloaded from Genbank with the R package *genomes* v2.12.0 (Stubben, 2014), and all data analysis was conducted in R with the following packages:ggplot2 v2.1.0, dplyr v0.4.3, tidyr v 0.4.1, and cowplot v0.6.2.

Further methodological details are provided in the *Supporting Information*.

## Acknowledgments

We thank Samuel Barnett for consultation on modeling the diffusive boundary layer effects and Chuck Pepe-Ranney for helpful discussions on the modeling approach used in this work. This material is based upon work supported by the Department of Energy, Office of Biological & Environmental Research Genomic Science Program under Award Numbers DE-SC0010558 and DE-SC0004486.

